# Functional Genomics Reveals TNT Bioremediation Strategies in *Pantoea* sp. MT58 and *Pseudomonas putida* KT2440

**DOI:** 10.64898/2026.04.16.711451

**Authors:** Audrey Liwen Wang, Thomas Eng, Alex Rivier, Saad Naseem, Alex Codik, Yan Chen, Aparajitha Srinivasan, Harshini Mukundan, Susannah G. Tringe, Christopher J. Petzold, Kara L. Nelson, Adam M. Deutschbauer, Aindrila Mukhopadhyay

## Abstract

2,4,6-Trinitrotoluene (TNT) is a recalcitrant and pervasive environmental pollutant. Although different environmental microbes have demonstrated their ability to degrade or transform TNT, the underlying genetic basis and cellular machinery remain unclear. In this study, we investigated bacterial strategies in response to TNT exposure in *Pantoea* sp. MT58 and *P. putida* KT2440 using proteomics and random barcode transposon-site sequencing (RB-TnSeq). *Pantoea* sp. MT58 was found to utilize TNT as a sole nitrogen source, whereas *P. putida* KT2440 exhibited only stress tolerance without assimilation. *Pantoea* sp. MT58 encodes multiple putative nitroreductases that were upregulated, yet deletion of these genes did not affect growth on TNT, revealing pathway redundancy. Furthermore, fitness profiling provided no evidence for genes involved in the canonical Meisenheimer-complex pathway associated with nitrite release. Instead, the data are most consistent with a sequential nitro-group reduction route in which nitrogen is ultimately recovered as ammonium, with nitrogen routed through the GS-GOGAT pathway with purine and urea pools as the candidate buffering architecture for TNT mineralization. Conversely, *P. putida* KT2440 relied on Ttg/RND efflux pumps and toluene tolerance proteins for survival without nitrogen assimilation from TNT. This work distinguishes routes for productive nitrogen assimilation from those involved in nitroaromatic tolerance, expanding the mechanistic understanding of anthropogenic compound metabolism to inform future bioremediation efforts.

## 1. Introduction

2,4,6-Trinitrotoluene (TNT) is a nitroaromatic energetic compound that persists in soils and groundwater at sites associated with mining,^1^ historical munitions production and military activities.^2,3^ Its extensive use throughout the twentieth century has resulted in widespread environmental dissemination, creating long-term contamination that continues to threaten ecological integrity and human health.^4,5^ The chemical structure of TNT, characterized by an electron-deficient aromatic ring with three symmetrically arranged nitro groups, makes it highly stable and recalcitrant to natural degradation processes, allowing it to persist in soil and groundwater for decades.^6^

Bioremediation offers an economically and environmentally feasible strategy for mitigating TNT contamination.^7^ While many TNT-degrading microbes have been isolated, biochemical data supporting its transformation has only been produced for limited pathways. One of the pathways involves the formation of a hydride-Meisenheimer complex as a key intermediate, which subsequently re-aromatizes TNT through the release of a nitrite ion (NO_2_^−^).^8–10^ Nitrite release is often a defining step of this pathway. The released nitrite is then reduced to ammonium (NH_4_^+^) and incorporated into central metabolism to support biomass generation.^8–10^ Microorganisms like *Pantoea* sp. Thu-Z and BJ2,^11,12^ *Enterobacter cloacae* PB2,^13^ *Pseudomonas putida* JLR11,^14^ and *Mycobacterium* sp.,^15^ have been reported to degrade TNT via a Meisenheimer-related pathway, sometimes solely based on the identification of homologous genes by a genome-level bioinformatic analysis. For example, a study^14^ using random transposon mutagenesis in *P. putida* JLR11 identified mutants defective in TNT mineralization and revealed that genes involved in nitrite catabolism were essential components of the TNT assimilation pathway. Importantly, the nitrite released during TNT degradation was further assimilated into central metabolism, supporting biomass production and cell viability.^14,16^

Another major TNT reduction pathway is through the sequential reduction of nitro groups to nitroso (−NO), hydroxylamino (HADNT), and amino derivatives (ADNT).^8–10^ Instead of nitrite release, nitrogen is liberated as ammonium (NH_4_^+^) from reactive intermediates, specifically HADNT derivatives.^8,17^ One proposed mechanism for this release is the Bamberger re-arrangement, where hydroxylamino intermediates yield aminophenols which subsequently release ammonia due to lyase activity or microbial aminophenol metabolism. Several microorganisms from divergent evolutionary clades, including *Escherichia coli*^18^ and *Pseudomonas* species^14,19–21^ are proposed to use this reduction pathway.

Apart from assimilating TNT as a nitrogen source, some bacteria also adopt a stress tolerance co-metabolism strategy by transforming TNT to mitigate toxicity without biomass gain.^22,23^ For example, *Stenotrophomonas* strain SG1 could transform roughly 32% of TNT, yet it failed to show significant growth unless supplemental carbon or nitrogen sources were provided.^23^

The strain *Pantoea* sp. MT58 was isolated from groundwater samples from the Oak Ridge Field Research site in Tennessee, USA.^24^ We hypothesize that the unique environmental stressors (low pH, high nitrate, heavy metal exposure) associated with this field site may select for novel metabolic adaptations that incidentally enable the utilization of this other anthropogenic molecule. Furthermore, other studies have implicated strains from the *Pantoea* genus in TNT tolerance and degradation.^11,12^ In this study, we applied an integrated multi-omics framework combining RB-TnSeq and proteomics to systematically investigate the mechanism of TNT metabolism in *Pantoea* sp. MT58, using the model organism *P. putida* KT2440 as a comparison. We demonstrate in this current study that *Pantoea* sp. MT58 could assimilate TNT as a nitrogen source, whereas *P. putida* KT2440 primarily adopted a stress-tolerance strategy to transform TNT.

## 2. Materials and Methods

### 2.1 Bacterial strains and growth media

All *Pantoea* sp. MT58 and *P. putida* KT2440 strains generated in this study are described in **Table S1**. Both strains were maintained on LB agar at 30 °C and stored as glycerol stocks at −80 °C. 2,4,6-Trinitrotoluene (TNT) was purchased as a 10 mg/mL solution in acetonitrile (MilliporeSigma Supelco, ERT107S5ML). TNT was recovered from the acetonitrile solvent by overnight evaporation. 1mL of the TNT acetonitrile solution was transferred to a microcentrifuge tube and incubated overnight in a room temperature fume hood. The recovered dried TNT was resuspended in 200 μL dimethyl sulfoxide (DMSO) and added to 100 mL M9 minimal salt medium (hereafter referred to as M9 medium) to achieve a final TNT concentration of 100 mg/L (0.44 mM, approximately 75% of saturation limit). The M9 medium was prepared following the NREL formulation^25^ with slight modifications that improve growth.^26^ M9 medium with DMSO is used for all matched control samples. For *Pantoea* sp. MT58 experiments using TNT as a sole nitrogen source, nitrogen salts (ammonium chloride, NH_4_Cl) were omitted from the M9 formulation.

### 2.2 Growth analysis using TNT as a nitrogen source

Growth curves were measured for both *Pantoea* sp. MT58 and *P. putida* KT2440 to assess TNT utilization as a sole nitrogen source. Cultures were grown in LB medium overnight and harvested by centrifugation (5,000 × *g*, 10 min), washed twice with NH_4_Cl-free M9 medium, and inoculated to an initial OD_600_ of 0.1 in the following three conditions: three biological replicates of M9 medium containing 1% glucose and 100 mM NH_4_Cl (positive control), three biological replicates of M9 medium with 1% glucose and 100 mg/L TNT (test condition), and one replicate of LB medium (rich medium control). Corresponding sterile medium blanks were included for each condition. Cultures were incubated at 30°C with orbital shaking (200 rpm) in 24-well microplates. Growth was monitored by measuring OD_600_ at 15-minute intervals over 30 hours using an automated microplate reader (Agilent Biotek Synergy H4).

### 2.3 Proteomics sample preparation and analysis

Overnight cultures of *Pantoea* sp. MT58 grown in LB medium were harvested by centrifugation (5,000 × *g*, 10 min, 4 °C) and then washed twice with nitrogen-free M9 medium to remove residual nitrogen sources. Cells were then resuspended in M9 medium containing 1% glucose supplemented with either 100 mg/L TNT (test condition) or 100 mM NH_4_Cl (control condition) as the sole nitrogen source. Cultures were incubated at 30 °C with shaking (200 rpm) and cells were harvested after 16 hours of growth by centrifugation.

Total protein was extracted from cell pellets and tryptic peptides were prepared by following established proteomic sample preparation protocol.^27^ Briefly, cell pellets were resuspended in Qiagen P2 Lysis Buffer (Qiagen, Germany) to promote cell lysis. Proteins were precipitated with addition of 1 mM NaCl and 4 x vol acetone, followed by two additional washes with 80% acetone in water. The recovered protein pellet was homogenized by pipetting mixing with 100 mM ammonium bicarbonate in 20% methanol. Protein concentration was determined by the DC protein assay (BioRad, USA). Protein reduction was accomplished using 5 mM tris 2-(carboxyethyl)phosphine (TCEP) for 30 min at room temperature, and alkylation was performed with 10 mM iodoacetamide (IAM; final concentration) for 30 min at room temperature in the dark. Overnight digestion with trypsin was accomplished with a 1:50 trypsin:total protein ratio. The resulting peptide samples were analyzed on an Agilent 1290 UHPLC system coupled to a Thermo Scientific Orbitrap Exploris 480 mass spectrometer for discovery proteomics.^28^ Briefly, peptide samples were loaded onto an Ascentis® ES-C18 Column (Sigma–Aldrich, USA) and were eluted from the column by using a 10 minute gradient from 98% solvent A (0.1 % FA in H_2_O) and 2% solvent B (0.1% FA in ACN) to 65% solvent A and 35% solvent B. Eluting peptides were introduced to the mass spectrometer operating in positive-ion mode and were measured in data-independent acquisition (DIA) mode with a duty cycle of 3 survey scans from m/z 380 to m/z 985 and 45 Tandem mass spectrometry (MS2) scans with precursor isolation width of 13.5 m/z to cover the mass range. DIA raw data files were analyzed by an integrated software suite DIA-NN.^29^ The databases used in the DIA-NN search (library-free mode) are *P. putida* KT2440 latest Uniprot proteome FASTA sequence and *Pantoea* sp. MT58 annotated proteome FASTA sequence from the fitness browser (https://fit.genomics.lbl.gov/cgi-bin/myFrontPage.cgi) plus the protein sequences of common proteomic contaminants. DIA-NN determines mass tolerances automatically based on first pass analysis of the samples with automated determination of optimal mass accuracies. The retention time extraction window was determined individually for all MS runs analyzed via the automated optimization procedure implemented in DIA-NN. Protein inference was enabled, and the quantification strategy was set to Robust LC = High Accuracy. The main DIA-NN output reports were filtered using a global false discovery rate (FDR) threshold of 0.01 (FDR ≤ 0.01) at both the precursor level and the protein group level. The Top3 method,^30^ which is the average MS signal response of the three most intense tryptic peptides of each identified protein, was used to plot the quantity of the targeted proteins in the samples. Differential abundance of proteins between test and control conditions was assessed using an unpaired two-tailed t-test. A significance threshold of p < 0.05 and fold-change (FC) ≥ 2 (|log_2_FC| ≥ 1) was applied to define differentially expressed proteins. FDR-adjusted p values were obtained using the Benjamini-Hochberg method,^31^ with results reported in **Figure S1**. Functional enrichment analysis of significant protein sets was performed using ShinyGO (v0.76).^32^

### 2.4. RB-TnSeq mutant library fitness assays and DNA sequencing

RB-TnSeq mutant libraries for *Pantoea* sp. MT58^24^ and *P. putida* KT2440^33^ were previously constructed following the protocol described by Wetmore et al.^34^ This procedure allowed the precise mapping of transposon insertion sites in relation to unique DNA barcodes. The *Pantoea* sp. MT58 library contains 447,153 uniquely mapped transposon mutants,^24^ and the *P. putida* KT2440 library contains approximately 185,401 mapped mutants.^26^ Frozen stocks of each mutant library were recovered in LB medium supplemented with 50 µg/mL kanamycin. *P. putida* KT2440 cultures were inoculated overnight, and *Pantoea* sp. MT58 cultures were inoculated in the morning; both libraries were harvested at the end of the day, washed twice with nitrogen-free M9 medium, and reference samples (time-zero) were collected. For mutant fitness experiments, the recovered libraries were inoculated into M9 medium at an initial OD_600_ of 0.05 under the specified test and control conditions.

For *Pantoea* sp. MT58, mutant fitness experiments were conducted with four biological replicates in M9 medium supplemented with 100 mg/L TNT as the sole nitrogen source, and three biological replicates in M9 medium supplemented with 100 mM NH_4_Cl as the control condition. For *P. putida* KT2440, the test conditions were four replicates of M9 with 100 mM NH_4_Cl and 100 mg/L TNT, and the control included a triplicate of M9 with 100 mM NH_4_Cl. The cells were harvested by centrifugation, and genomic DNA was extracted using the Wizard® Genomic DNA Purification Kit (A1120) (Promega Corporation, USA) following the manufacturer’s protocol.^35^ The extracted DNA samples were stored at −80 °C until use. Library preparation for barcode sequencing was performed using the BarSeq98 method,^34^ and the resulting PCR products were sequenced on an Illumina NovaSeq system (Illumina Inc., USA). Gene fitness values were calculated by comparing barcode abundances between time zero and post-exposure samples as previously described.^34^

In this study, mutant fitness values were calculated as log_2_ ratios representing the change in abundance of mutants associated with each gene between the time-zero reference sample and the final sample collected after growth under the specified conditions.^34^ Fitness values less than −1 were interpreted as indicating that disruption of the gene caused a substantial growth defect under the test condition, with increasingly negative values (e.g., −2, −3) corresponding to progressively larger fitness costs and greater importance of the gene for growth under that condition. In contrast, fitness scores greater than 1 indicated that gene disruption conferred a substantial growth advantage, while fitness between −1 and 1 reflected a subtle phenotype with little to no measurable effect on cell growth.

### 2.5. Single and double deletion of oxidoreductases using the SacB-based plasmid integration system

To evaluate gene essentiality in TNT degradation, single and double deletions of oxidoreductase genes IAI47_09985 and IAI47_07670 were generated in *Pantoea* sp. MT58 using an allelic exchange protocol with *sacB* counter-selection and R6K-origin plasmids (**Table S1** and **Table S2**). The genotyping primers are provided in **Table S3**. Briefly, deletion plasmids containing upstream and downstream 1000-bp homology regions flanking the target genes were first introduced into the conjugation donor strain *E. coli* WM3064 (*pir gene*+, diaminopimelic acid [DAP]-) by electroporation. Then, the WM3064 donor cells were washed twice with 10% glycerol, conjugated with the recipient *Pantoea* sp. MT58 cells at a 1:1 ratio by incubating overnight on LB+ 300 µM DAP plates at 30 °C. Transconjugants were selected on LB+kanamycin plates, confirming successful plasmid integration into the *Pantoea* sp. MT58 chromosome.

Integrated strains were grown overnight in non-selective LB medium to promote spontaneous plasmid excision events. Then, cultures were plated on LB medium containing 5% sucrose and incubated at room temperature for two days to select for sacB counter-selection events. Resultant colonies were screened by patch testing on both kanamycin- and sucrose-containing media, with candidates that grew on sucrose but not on antibiotic medium subjected to PCR verification to confirm successful gene deletion. Double deletion mutants were generated by sequential application of this protocol to single deletion strains. For growth analysis of the oxidoreductase mutants, wild-type *Pantoea* sp. MT58 and single/double-deletion mutants were cultured in M9 medium supplemented with a nitrogen source in the form of either 100 mM NH_4_Cl or 100 mg/L 2,4,6-TNT. Growth curves were monitored by measuring OD_600_ over time. Bacterial viability was assessed through colony-forming unit (CFU) enumeration using serial 10-fold dilutions plated on LB selective media. Genomic DNA was extracted using a Wizard® genomic DNA purification kit (A1120) (Promega Corporation, USA) and Oxford Nanopore long read sequencing technology was used for whole genome sequencing (Plasmidsaurus Inc, USA). Raw sequencing data was mapped to the reference genome using Geneious Prime (v.2026.0) and the anticipated mutations were manually inspected for further clarification.

### 2.6 HPLC quantification of TNT and its intermediates in *P. putida* KT2440

Although *P. putida* KT2440 was unable to use TNT as its sole nitrogen source in M9 medium, we hypothesized that TNT transformation might still occur. Samples were prepared for HPLC analysis to detect TNT and its intermediates. The sample set consisted of *P. putida* KT2440 cultures grown for 24 hours in M9 medium supplemented with 100 mM NH_4_Cl and 100 mg/L TNT, as well as corresponding media-only controls containing the same supplements but no *P. putida* KT2440. Samples were first centrifuged at 4,000 rpm for 10 min at room temperature to remove biomass. 200 µL of the resulting supernatant were transferred to a 0.2 µm filter plate (Corning, USA) and centrifuged at 4,500 rpm at room temperature for 5 min to remove particulates. All measurements were performed on an Agilent 1260 Infinity Series HPLC system (Agilent Technologies, USA) equipped with 1260 series quaternary pump (G7104C), multi sampler (G7167A), thermostatted column compartment (G7116A), and diode array detector (G7117C). HPLC analyses were carried out with Inertsil diol (4.6 mm×150 mm, 3 µm) as described in a previous study.^36^ Briefly, acetonitrile-water gradient was used as the mobile phase with the initial acetonitrile:water ratio raised from 10% to 60% in 2–20 min and a final gradual reduction to 10% in a total runtime of 25 min at 0.4 mL/min. For all measurements, room temperature was used as the system temperature. Sample injection volume was 5 µL and the detection was carried out using the DAD detector at 254 nm. The data was analyzed using the OpenLab CDS v2.3.53 (Agilent Technologies, USA). Peaks were identified based on retention times relative to TNT standards prepared in DMSO and diluted serially in acetonitrile.

### 2.7 Phylogenetic analysis to identify potential oxidoreductase/nitroreductase homologs

Candidate oxidoreductase and nitroreductase protein sequences from *Pantoea* sp. MT58 and *P. putida* KT2440 were analyzed to identify homologs of previously characterized TNT-degrading or nitroaromatic reducing enzymes. Sequence information on previously characterized TNT enzymes is provided in **Table S4**, and their protein sequences were downloaded from NCBI. Multiple sequence alignment was performed using MUSCLE with default parameters. A phylogenetic tree was constructed in Geneious Prime (v.2026.0) using the neighbor-joining method with the Jukes-Cantor distance model. Node support was assessed with 1,000 bootstrap replicates, and a 40% support threshold was applied to generate a consensus tree. The human NAD(P)H:quinone oxidoreductase 1 (NQO1) sequence was included as an outgroup to root the tree.

## 3. Results

### 3.1 *Pantoea* sp. MT58 can utilize TNT as a sole nitrogen source

To investigate potential TNT metabolism in *Pantoea* sp. MT58, we first assessed its ability to grow on TNT as the sole nitrogen source. *Pantoea* sp. MT58 exhibited robust growth in M9 medium supplemented with 100 mg/L TNT as the sole nitrogen source (**Figure 1A**, red line). Although the growth rate and final cell density were lower than those observed with 100mM NH_4_Cl as the nitrogen source (light gray line) or in LB medium (dark gray line), due to the lower concentration of TNT supplied, *Pantoea* sp. MT58 still demonstrated the ability to use TNT to support biomass production without additional nitrogen supplementation.

**Figure 1:**
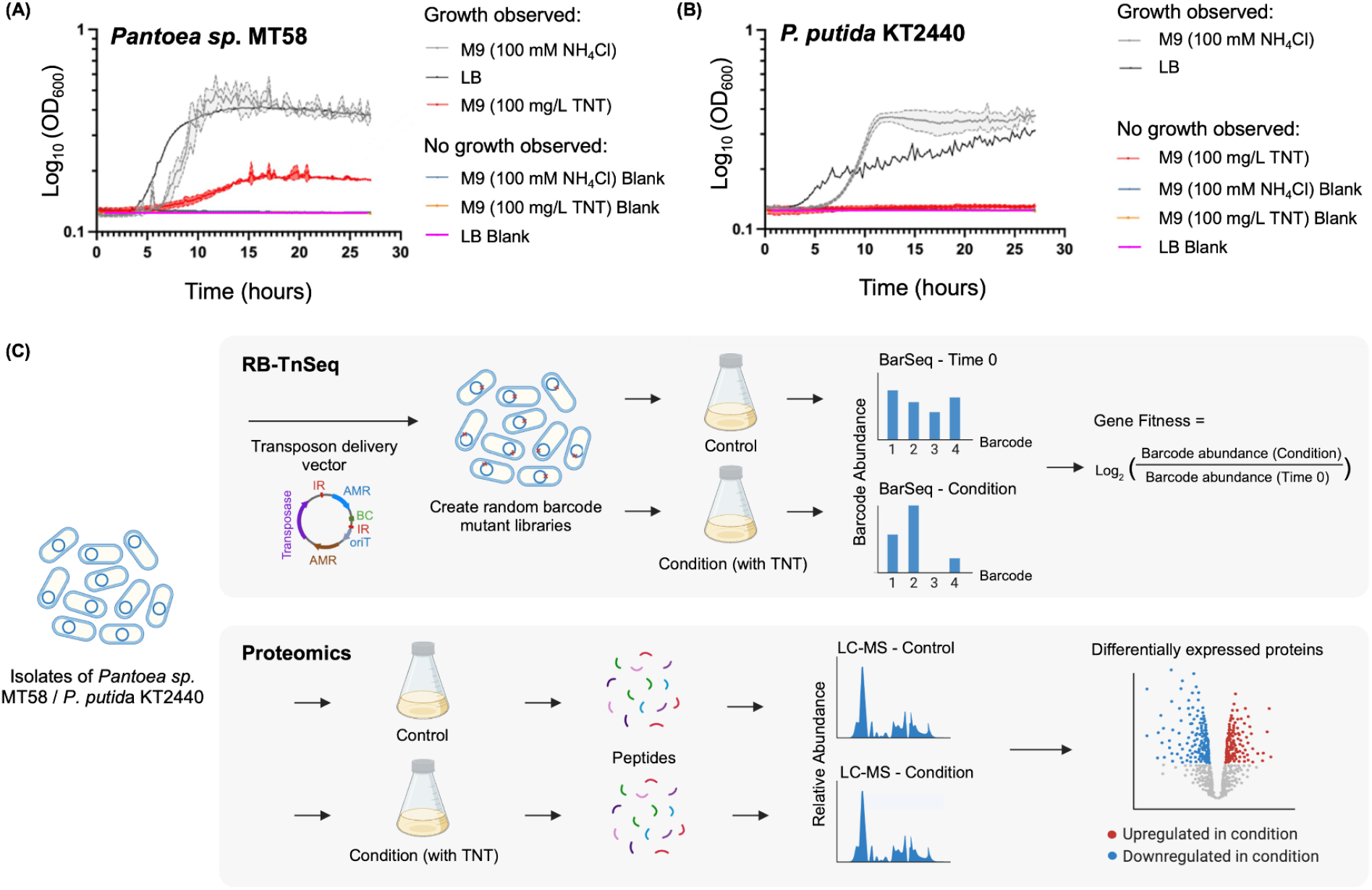
Growth curves for **(A)** *Pantoea* sp. MT58 and **(B)** *P. putida* KT2440 cultured in M9 medium supplemented with 100 mM NH_4_Cl (light gray) or 100 mg/L TNT (red), as well as in LB medium (dark gray). Media-only blank controls were included for each condition (NH_4_Cl, blue; TNT, orange; LB, pink). **(C)** Schematic of experimental design for functional genomics approaches (RB-TnSeq and proteomics). The top panel illustrates the RB-TnSeq workflow. In this approach, random barcode mutant libraries were created previously for both *Pantoea* sp. MT58^24^ and *P. putida* KT2440^38^, followed by competitive mutant fitness assays. These assays involved comparing the abundance of DNA barcodes using BarSeq before (Time 0) and after selective growth under TNT conditions. The comparison allows for the calculation of gene fitness by assessing how the relative abundance of each barcoded transposon mutant changes in response to the selective pressure of TNT, providing insights into the phenotypes and fitness associated with specific gene mutations. The bottom panel depicts the proteomics approach, where isolates are grown in parallel under control and TNT-supplemented conditions. Following growth, samples undergo enzymatic digestion to prepare peptides for LC-MS analysis. Differentially expressed proteins between the control and TNT-supplemented conditions are identified. Figure C panels generated using Biorender.

For comparison, we tested the well-characterized model organism *P. putida* KT2440, which is not a known TNT specialist, under the same conditions. We were initially motivated to determine if *P. putida* KT2440 could also assimilate TNT due to its high average nucleotide identity to a known TNT assimilator, *P. putida* JLR11. However, the genome assembly of JLR11 does not currently contain a recircularized genome and only has 39 contigs with low sequence coverage,^37^ making an accurate comparison of genome content challenging. *P. putida* KT2440 grew robustly in M9 medium supplemented with NH_4_Cl (**Figure 1B**, light gray line) and in LB medium (dark gray line); however, it failed to grow when TNT was the sole nitrogen source (red line), suggesting that TNT was imposing stress rather than supporting growth for the strain. This result confirms that the ability to use nitrogen from TNT is a specialized metabolic trait not present in the standard *P. putida* KT2440 laboratory strain. As *P. putida* KT2440 was unable to use TNT as a sole nitrogen source, we categorized its response as co-metabolizing (M9 medium with both NH_4_Cl and TNT) for the multi-omics experiments.

### 3.2 A multi-omics approach to elucidate bacterial strategies under TNT exposure

We applied a multi-omics approach to investigate the genetic basis of TNT use in *Pantoea* sp. MT58 and TNT stress tolerance in *P. putida* KT2440. First, we performed a randomly barcoded transposon sequencing (RB-TnSeq) experiment to identify genes whose functions were important for growth or survival on TNT (**Figure 1C**). The pooled mutant libraries for each strain contained over a hundred thousand unique insertion mutants and were grown competitively in control and TNT-containing conditions. The relative fitness of each gene knockout was then calculated by quantifying the change in its corresponding barcode abundance via high-throughput sequencing (BarSeq). This functional genomics approach allows for the identification of genes that are important for fitness, as assessed by single-gene mutations.

Second, to identify the cellular machinery that is actively deployed in response to TNT, we performed quantitative proteomics (**Figure 1C**). Both strains were cultured in both control and TNT-containing M9 medium, and the relative abundances of approximately 3,000 proteins were determined by LC-MS. Proteomics enabled the identification of differentially expressed proteins, revealing pathways and systems that were upregulated in response to TNT catabolism and stress, as well as those that were downregulated to conserve cellular resources. Together, the RB-TnSeq and proteomics provided a systems-level view of the metabolic and regulatory logic underlying TNT use in *Pantoea* sp. MT58 and stress tolerance in *P. putida* KT2440.

### 3.3 Stress tolerance of *P. putida* KT2440 in response to TNT exposure

For *P. putida* KT2440, proteomics data revealed significant changes in protein abundance under TNT exposure, with 100 proteins significantly upregulated and 65 proteins significantly downregulated relative to the control condition (**Figure 2A**). Functional enrichment analysis of the upregulated proteins showed a strong signature of secondary metabolite biosynthesis and stress response regulation. The most significantly enriched terms were associated with nonribosomal peptide synthetase (NRPS) and polyketide synthase (PKS) machinery, including condensation domains, phosphopantetheine attachment sites/binding domains, and amino acid adenylation domains (**Figure 2B**), which are all important components for assembling complex bioactive peptides and polyketides.^39,40^ Additionally, the enrichment of translational regulator CsrA and the carbon storage regulator superfamily indicates activation of the Gac/Rsm post-transcriptional regulatory cascade, which globally controls secondary metabolism^41,42^ (**Figure 2B**). Additional enrichment associated with ACP-like superfamily, oxidoreductase activity, and oxidation-reduction process were observed (**Figure 2B**), indicating stress response regulation.^43^ However, there was no significant enrichment related to nitrogen assimilation pathways.

**Figure 2:**
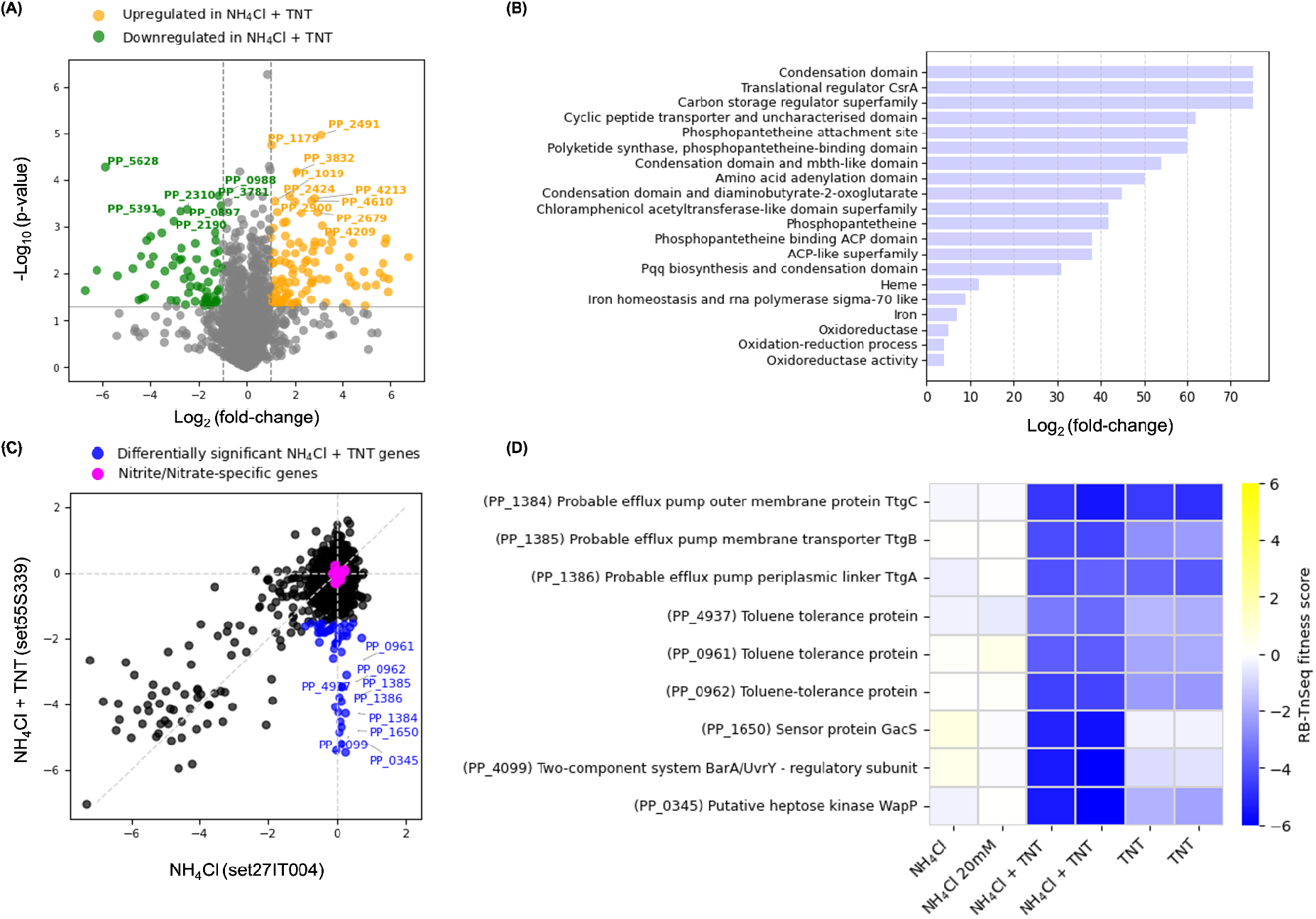
Stress tolerance of *P. putida* KT2440 in response to TNT exposure. **(A)** Proteomics data highlighting differentially expressed proteins when comparing growth in NH_4_Cl versus NH_4_Cl + TNT. The scatter plot shows log_2_ fold change against the negative log_10_ p-value for each protein, with proteins exhibiting a fold change of ±2 and a p-value < 0.05 highlighted in orange (upregulated) and green (downregulated) in the NH_4_Cl + TNT condition. **(B)** Fold enrichment of differentially significant genes in NH_4_Cl + TNT analyzed by ShinyGo, with the bars illustrating the fold enrichment of mapped gene functions. **(C)** Scatter plot comparing gene fitness scores under control NH_4_Cl and NH_4_Cl+TNT conditions. Differentially significant genes are highlighted in blue, indicating genes with near-neutral fitness in NH_4_Cl alone (−1 ≤ fitness ≤ 1) and a strong fitness defect under NH_4_Cl + TNT conditions (fitness < −1.5). Nitrite/nitrate-specific genes are indicated in pink. **(D)** Heatmap displaying the fitness scores of significant genes under three conditions: NH_4_Cl only, NH_4_Cl + TNT, and TNT only. Even though TNT alone did not support measurable growth, we still included the Tn-Seq results to assess gene fitness under survival or persistence conditions. Repeated columns are replicates.

The stress-response signature of *P. putida* KT2440 was further clarified by the RB-TnSeq fitness data. A comparison of gene fitness between the control condition and the TNT co-metabolizing condition indicated that 47 genes were important for *P. putida* KT2440’s tolerance under TNT exposure (**Figure 2C**, blue dots). A heatmap of these genes highlights the critical roles of the toluene tolerance proteins (PP_4937, PP_0961, and PP_0962) and the associated efflux pump system (TtgABC, PP_1384-PP_1386), as well as a sensor protein (PP_1650) and a two-component regulatory system (PP_4099) known to be involved in stress responses^26,44^ (**Figure 2D**). The genes specific for nitrite and nitrate metabolism were not important for *P. putida* KT2440 under either condition (**Figure 2C**, pink dots). Although there was no evidence of TNT-derived nitrogen assimilation in *P. putida* KT2440, HPLC measurements showed that TNT concentration decreased to below the detection limit after 24 h (**Table S5**).

We further investigated oxidoreductases potentially involved in TNT transformation. An oxidoreductase (PP_4931, ~2.3-fold) and an azoreductase (PP_2866, ~11.1-fold) were significantly upregulated under TNT conditions based on the proteomics data (**Figure 2A, Table S6**). Phylogenetic analysis indicated that PP_2866 is a close homolog (70.4% identity) of an azoreductase in *Pseudomonas aeruginosa* involved in TNT degradation (**Figure 4A**). RB-TnSeq data suggest that PP_4931 is essential for viability, as no corresponding mutants were recovered, while PP_2866 did not exhibit a significant fitness phenotype (**Table S7**). Collectively, these results indicate that *P. putida* KT2440 could partially transform TNT to mitigate the toxicity but could not utilize it as a nitrogen source.

### 3.4. Functional redundancy of oxidoreductases in *Pantoea* sp. MT58

To investigate how *Pantoea* sp. MT58 utilizes TNT as a nitrogen source, we examined nitroreductases/oxidoreductases potentially capable of reducing nitro-groups on aromatic compounds. From the proteomics data, we observed that multiple nitroreductases or oxidoreductases were significantly upregulated in *Pantoea* sp. MT58 during growth in TNT (**Figure 3A**). Two NADP(H)-dependent oxidoreductases were upregulated, including a short-chain dehydrogenases (SDR) family enzyme (IAI47_07670), which exhibited the highest fold-change (~21-fold), and another oxidoreductase (IAI47_09985, ~4-fold). Both subunits of xanthine dehydrogenase (IAI47_09815 and IAI47_09820) showed greater than a 10-fold differential increase. A nitroreductase family protein (IAI47_19540) was upregulated (~3.6-fold), which may initiate reduction of TNT nitro groups. Increased protein levels were also observed for an alkene reductase (IAI47_10765, ~5.7-fold) and another SDR family oxidoreductase (IAI47_01395, ~3.9-fold). However, when we examined the genes for these proteins in the RB-TnSeq dataset, we did not find corresponding fitness changes from the TNT fitness growth assay (**Figure 3B**, black-labeled proteins). We hypothesized that there might be functional redundancy in these oxidoreductases so that single-gene deletion did not lead to fitness defects.

**Figure 3.**
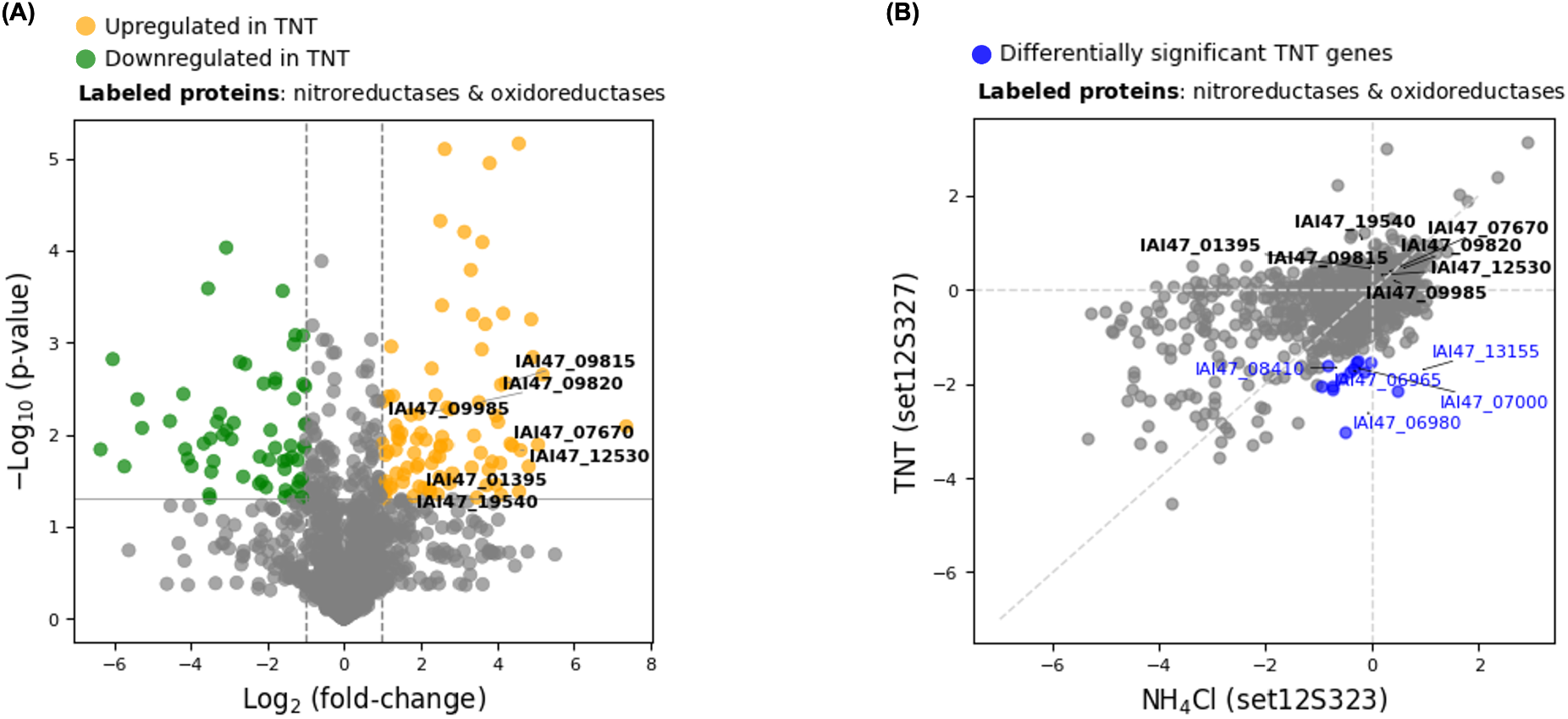
Upregulation of nitroreductases/oxidoreductases in proteomic data without corresponding fitness changes in RB-TnSeq for *Pantoea* sp. MT58. **(A)** Proteomics data highlighting differentially expressed proteins when comparing growth in 100 mM NH_4_Cl versus 100 mg/L TNT as the sole nitrogen source in M9 medium. The scatter plot shows log_2_ fold change against the negative log_10_ p-value for each protein, with proteins exhibiting a fold change of ±2 and a p-value < 0.05 highlighted in orange (upregulated) and green (downregulated) in the TNT condition. Seven nitroreductases and oxidoreductases potentially involved in the initial breakdown of TNT are labeled in black. **(B)** Scatter plot comparing gene fitness scores from RB-TnSeq data between control (NH_4_Cl) and TNT conditions. Differentially significant genes under TNT condition are colored in blue, and the same set of nitroreductases and oxidoreductases are labeled in black.

Both NADP-related oxidoreductases (IAI47_07670 and IAI47_09985) were of particular interest due to the fact that many nitroaromatic-transforming oxidoreductases rely on NADPH as an electron donor in reduction reactions. IAI47_07670 was also the most highly upregulated oxidoreductase in our proteomics dataset (~21-fold). Therefore, we hypothesized that these NADP-related enzymes were the candidate contributors to an early reductive step in TNT assimilation. To test this, we constructed single (ΔIAI47_07670, ΔIAI47_09985) and double (ΔIAI47_07670 + ΔIAI47_09985) deletion mutants.

We first confirmed that the deletion of these genes had no significant impact on general fitness by growing the wild-type and mutant strains in M9 medium with NH_4_Cl as the nitrogen source. All strains exhibited nearly identical growth curves, reaching the same final cell density (**Figure 4B**). Subsequently, we assessed the growth of these strains in M9 medium with 100 mg/L TNT as the sole nitrogen source. Neither the single deletion mutants nor the double deletion mutant showed a significant growth defect compared to the wild-type strain (**Figure 4C**). All strains were able to utilize TNT as a nitrogen source, reaching similar final cell densities as the WT after approximately 20 hours. To confirm that the observed optical density corresponded to viable cells, we performed a 10-fold serial dilution and spotting assay from the liquid cultures after 25 hours. This assay confirmed that all four strains, including wild-type, the two single mutants, and the double mutant, achieved comparable, high cell densities of approximately 8 x 108 CFUs/mL (**Figure 4D**).

**Figure 4:**
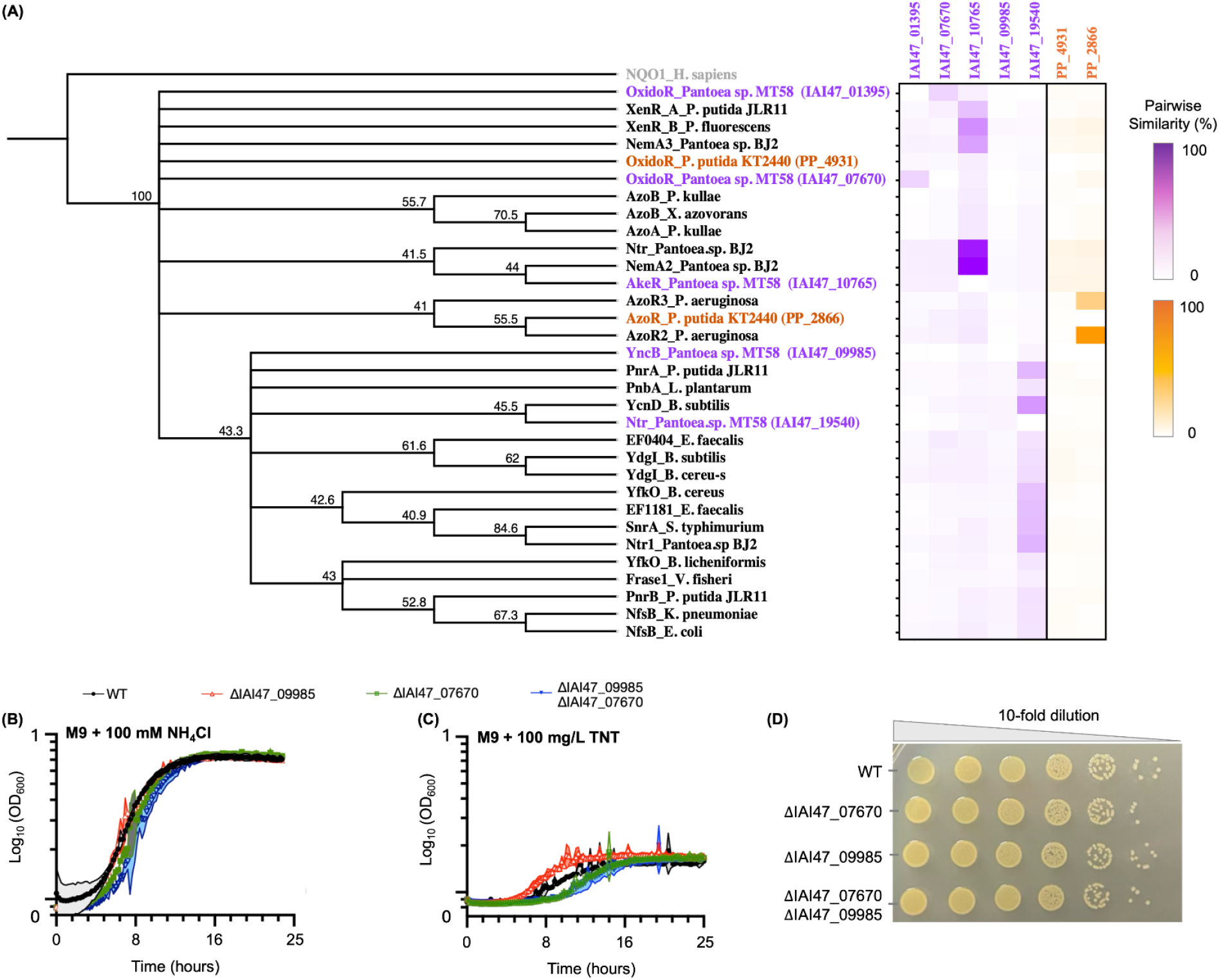
Functional redundancy of enzymes responsible for TNT reduction in *Pantoea* sp. MT58. **(A)** A phylogenetic tree was constructed to assess whether oxidoreductases/nitroreductases identified in *Pantoea* sp. MT58 and *P. putida* KT2440 are homologous to previously characterized TNT-degrading or xenobiotic-metabolizing enzymes. Candidate proteins from *P. putida* KT2440 are highlighted in orange, and those from *Pantoea* sp. MT58 in purple. Reference protein sequences were obtained from NCBI (**Table S4**). The tree was generated as described in **Section 2.7**, with bootstrap support values (%) indicated at the nodes. NQO1 was used as an outgroup. Pairwise distance (%) derived from the phylogenetic tree was used to generate the heatmap shown on the right. Color intensity reflects sequence similarity (%), with darker colors indicating higher similarity. **(B-D)** Growth analyses of wild-type (WT) and nitroreductase mutants (ΔIAI47_09985, ΔIAI47_07670, and ΔIAI47_09985 + ΔIAI47_07670) in *Pantoea* sp. MT58. Growth curves for the single and double deletion mutants, as well as the wild type cultured in M9 medium supplemented with **(B)** 100 mM NH_4_Cl and **(C)** 100 mg/L TNT. **(D)** Colony-forming units (CFUs) obtained from a 10-fold dilution series for each mutant type and the wild type after growth in M9 TNT media.

The finding that simultaneous deletion of the two most highly upregulated nitroreductase candidates had no measurable impact on the ability of *Pantoea* sp. MT58 to assimilate TNT suggests functional redundancy in the initial reductive steps or the involvement of an essential gene that is not represented in the RB-TnSeq dataset. We further investigated additional candidate proteins by performing phylogenetic analysis and examining sequence similarity to identify potential homologs of previously characterized TNT-degrading or nitroaromatic-reducing enzymes (**Figure 4A**). Based on the similarity heatmap in **Figure 4A**, an alkene reductase (IAI47_10765) from *Pantoea* sp. MT58 exhibited 89% sequence similarity to a nitroreductase and 98% similarity to an N-ethylmaleimide reductase, both previously involved in TNT metabolism in *Pantoea* sp. BJ2.^12^ In addition, IAI47_10765 shared 44.7% sequence similarity with xenobiotic reductase B from *P. fluorescens*,^45^ which is associated with TNT degradation. A nitroreductase (IAI47_19540) from *Pantoea* sp. MT58 showed 29.3% similarity to the TNT-degrading PnrA nitroreductase from *P. putida* JLR11^46^ and 41% similarity to the nitro-reducing YcnD oxidoreductase from *Bacillus subtilis* ^47^(**Figure 4A**).

### 3.5 A complementary urea-related nitrogen assimilation pathway in *Pantoea* sp. MT58

The involvement of nitrite- and nitrate-related genes in the canonical Meisenheimer-complex pathway provides evidence for the pathway’s role in TNT transformation. In contrast, their absence suggests a sequential nitro-reduction pathway, in which nitrogen may instead be released as ammonium.^8–10^ To understand the nitrogen assimilation process in *Pantoea* sp. MT58, we examined differential RB-TnSeq fitness profiles of genes across multiple nitrogen sources, including nitrate, nitrite, NH_4_Cl, and TNT. While genes for core biosynthetic pathways like purine/pyrimidine nucleotide synthesis were important under all nitrogen-limiting conditions (top box, **Figure 5A**), a clear separation emerged for the nitrogen-processing machinery (middle box, **Figure 5A**). Genes annotated as nitrite/nitrate sensing proteins and reductases, were highly important for growth on nitrate and nitrite, but showed no fitness defect when TNT or ammonium was the nitrogen source. This suggests that nitrogen from TNT is likely not released as nitrite in *Pantoea* sp. MT58 but instead as ammonium, supporting a sequential nitro-reduction pathway rather than the Meisenheimer-complex pathway. Additionally, although TNT can serve as a nitrogen source, it also imposes stress on *Pantoea* sp. MT58, as genes related to lipid membrane stress were found to be important (bottom box, **Figure 5A**).

**Figure 5:**
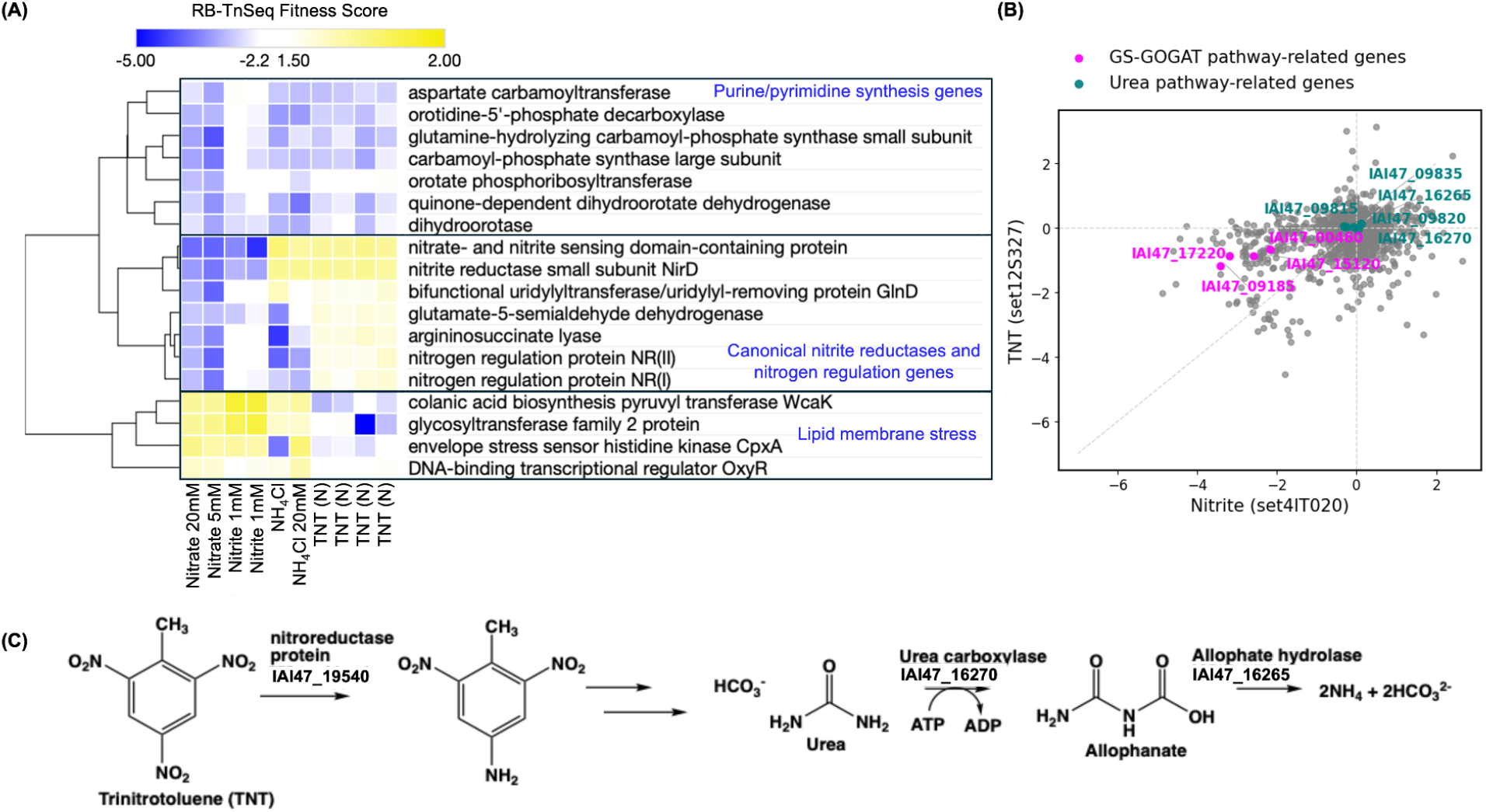
Proposed urea-centric nitrogen assimilation pathway in *Pantoea* sp. MT58. **(A)** Heatmap of RB-TnSeq fitness scores for *Pantoea* sp. MT58 genes under different nitrogen sources: nitrate (5 mM, 20 mM), nitrite (1 mM, two replicates), NH_4_Cl (20 mM, two replicates), and TNT (100 mg/L, four replicates). The blue-to-yellow color gradient represents fitness scores ranging from negative (gene is important under the condition) to positive (gene is costly under the condition). Highlighted categories include genes involved in purine/pyrimidine synthesis (top box), canonical nitrite assimilation (middle box), and lipid membrane stress responses (bottom box). **(B)** Scatter plot comparing gene fitness scores under nitrite and TNT conditions. Genes associated with the GS-GOGAT pathway are highlighted in pink, and those associated with the urea pathway are highlighted in teal. **(C)** Proposed nitrogen assimilation pathway in *Pantoea* sp. MT58, including enzymes like nitroreductase (IAI47_19540), urea carboxylase (IAI47_16270), and allophanate hydrolase (IAI47_16265).

In parallel, proteomics revealed a significant upregulation of urea-related genes, indicating activation of a urea import and catabolism pathway. Based on this, we propose a putative urea-centric nitrogen assimilation route (**Figure 5C**). In this pathway, xanthine dehydrogenase subunits (IAI47_09815, ~8.2-fold; IAI47_09820, ~7-fold) and urate hydroxylase (IAI47_09835, ~8.2-fold) convert purines through intermediates such as xanthine and uric acid, yielding urea. The urea is then processed by urea carboxylase (IAI47_16270, ~10.1-fold) to form allophanate, which is subsequently hydrolyzed by allophanate hydrolase (IAI47_16265, ~7.2-fold), releasing two molecules of ammonia. This upregulated pathway exhibited neutral fitness scores (fitness ≈ 0) when assayed on TNT (**Figure 5B**, teal dots). In contrast, genes involved in the canonical GS-GOGAT nitrogen assimilation pathway showed fitness defects under both TNT and nitrite conditions (**Figure 5B**, pink dots). This suggests that GS-GOGAT remains the primary route for nitrogen assimilation in *Pantoea* sp. MT58. The urea pathway, while transcriptionally upregulated, may therefore play a complementary role in nitrogen utilization.

## 4. Discussion

### 4.1 Productive nitrogen assimilation versus stress tolerance strategy under TNT exposure

The combined use of proteomics and genome-wide gene fitness profiling reveals two bacterial strategies for responding to TNT: assimilation of TNT-derived nitrogen to support growth versus stress tolerance without growth. A notable finding in this study is the putative functional redundancy in *Pantoea* sp. MT58’s TNT degradation machinery. We propose that *Pantoea* sp. MT58 adopts a generalist strategy for degrading anthropogenic compounds. Potentially by the selection pressures inherent to the Oak Ridge Field research site groundwater and sediment, *Pantoea* sp. MT58 constitutively expresses a set of broad-spectrum nonspecific oxidoreductases, while inducing a diverse set of nitroreductases upon TNT exposure.

In the Meisenheimer-complex pathway for aerobic TNT degradation, genes encoding nitrite reductases should be essential for growth on TNT, as cells must detoxify the nitrite flux to survive.^8–10^ This pathway has been reported in *Pantoea sp*. Thu-Z, which assimilates up to 96.6% of TNT-derived nitrogen into biomass through direct aromatic ring reduction.^11^ Another strain *Pantoea sp*. BJ2 shows strong evidence of stepwise reduction of nitro groups, although proteins involved in nitrate/nitrite transport and reduction are also significantly upregulated.^12^ However, our functional genomics data provide evidence against the Meisenheimer-complex pathway in *Pantoea* sp. MT58 and instead support a sequential nitro-reduction pathway. Consistent with this, an alkene reductase (IAI47_10765) in *Pantoea sp*. MT58 shares up to 98% sequence similarity with nitroreductases (Ntr and NemA2) in *Pantoea sp*. BJ2, which are responsible for reducing nitro groups on the TNT aromatic ring to hydroxylamino or amino derivatives.

On the other hand, *P. putida* KT2440 transformed TNT, followed by efflux as part of a well-characterized detoxification mechanism.^48^ An azoreductase (PP_2866) in *P. putida* KT2440 showed high sequence similarity to a previously characterized TNT-transforming azoreductase in *P. aeruginosa*,^49,50^ which may explain the complete transformation of TNT within 24 h. Despite efficient transformation and subsequent efflux, no bacterial growth was observed in our study, even though previous studies showed certain *Pseudomonas* strains could utilize TNT as a sole nitrogen source.^14,37,49–51^

### 4.2 A nitrogen-buffering architecture for *Pantoea* sp. *MT58*

Proteomics revealed substantial upregulation of the urea ABC transporter (UrtABCDE), urea-degrading enzymes urea carboxylase (IAI47_16270), and allophanate hydrolase (IAI47_16265). Nevertheless, RB-TnSeq showed that these urea-processing genes were dispensable for survival on TNT. In contrast, genes for purine and pyrimidine biosynthesis exhibited strong negative fitness scores, indicating they are important for growth with TNT as the sole nitrogen source. The above evidence points to a model where nucleotide metabolism acts as the primary nitrogen sink. TNT-derived nitrogen is assimilated into glutamine and glutamate via the GS-GOGAT pathway, which act as central nitrogen donors for nucleotide biosynthesis.^52,53^

The urea pathway likely functions as a high-capacity overflow valve or nitrogen capacitor. When the primary nucleotide sink approaches saturation, excess nitrogen is shunted through catabolic nodes via purine degradation to generate urea. This urea could then be recycled to ammonium via the upregulated urea carboxylase system and assimilated back through the GS-GOGAT pathway.^54–58^ This buffering architecture may allow the cell to uncouple nitrogen assimilation from immediate growth demand, maintaining steady and controlled nitrogen availability under variable TNT flux.

### 4.3 Toward effective TNT bioremediation: implications and limitations

The ability of *Pantoea* sp. MT58 to utilize TNT as a nitrogen source during its degradation suggests a more self-sustaining bioremediation strategy compared to organisms that rely on co-metabolism, particularly under nitrogen-limited conditions. Moreover, the presence of nonspecific oxidoreductases and functional redundancy indicates that TNT reduction in *Pantoea* sp. MT58 is more metabolically robust and likely resilient to fluctuating environmental conditions. Such redundancy may confer an advantage for bioremediation systems, where microbial communities could experience variable substrate availability or shifting redox gradients.^59,60^ However, microbial TNT transformation or degradation often does not result in complete mineralization to non-toxic end products. In most characterized systems, nitroreductase activity often leads to the accumulation of ADNT and DANT isomers as terminal or near-terminal products. These compounds remain toxic and mutagenic, and are often resistant to further degradation.^7,9^ Therefore, understanding the metabolic fate of TNT and its intermediates is critical for sustainable bioremediation applications.

## 5. Acknowledgements

Funding for this project was from the Defense Advanced Research Projects Agency (DARPA) under contract O2503-097-089-091224 with Lawrence Berkeley National Laboratory for the PEDOMETER (Plant-Enhanced Degradation Of Munitions by Engineered TERrestrial microbes) project (ended Oct 2025). ALW performed work as a visiting researcher at LBNL. We thank Romy Chakraborty for helpful discussions and team membership on the PEDOMETER project.

## 6. Data availability

The RB-TnSeq datasets will be publicly available upon publication and can be accessed from the Fitness Browser (https://fit.genomics.lbl.gov/cgi-bin/myFrontPage.cgi). Fitness Browser dataset identifiers for *Pantoea* sp. MT58 includes set12S327 through set12S330 for the TNT condition and set12S323, set4IT017, and set4IT022 for the control fitness conditions. Datasets for *Pantoea* sp. MT58 grown with nitrite or nitrate are available under set4IT020 and set4IT023 through set4IT025. For *P. putida* KT2440, dataset identifiers include set55S339 through set55S342 for the TNT condition and set27IT004 through set27IT005 for the control fitness condition. Whole genome sequencing data for the *Pantoea* sp. MT58 single and double deletion mutants have been deposited at NCBI SRA under BioProject ID PRJNA1420862 with the locus tag prefix ACZB7N and will be released upon publication of this manuscript. The mass spectrometry proteomics data have been deposited to the ProteomeXchange Consortium via the PRIDE partner repository^61^ with the dataset identifier PXD073692. DIA-NN is freely available for download from https://github.com/vdemichev/DiaNN. Information on the strains and plasmids is provided in **Table S1** and **Table S2**. Strains and plasmids are available from the corresponding authors upon request, subject to review and completion of a material transfer agreement. Raw protein counts are provided in **Table S6**. Requests should be directed to.

