## Supplementary Tables and Figures for "Functional Genomics Reveals TNT Bioremediation Strategies in *Pantoea* sp. MT58 and *Pseudomonas putida* KT2440"

Number of Pages: 8
Number of Tables: 7
Number of Figures: 1

#### Table of Contents

**Supplementary Figures**

**Figure S1.** Adjusted p-value proteomics data of (A) *P. putida* KT2440 and (B) *Pantoea* sp. MT58

**Supplementary Tables**

**Table S1.** Strains used in this study
**Table S2.** Plasmids used in this study
**Table S3.** Genotyping primers
**Table S4.** Reference sequences for plotting the phylogenetic protein tree
**Table S5.** Residual TNT concentration (mg/L) in P. putida KT2440 cultures at 0 and 24 h
**Table S6.** Proteomics data of **(A)** *P. putida* KT2440 and **(B)** *Pantoea sp.* MT58. The data are provided as a separate Excel file Supplementary Table S6.
**Table S7.** RB-TnSeq gene fitness scores for an oxidoreductase (PP_4931) and an azoreductase (PP_2866)

#### Supplementary Tables

**Table S1.** Strains used in this study.

| Strain | Description | Reference |
|---|---|---|
| *P. putida* KT2440 | *Pseudomonas putida* KT2440 prototroph CmR AmpR | JBEI-13809 ATCC 47054 [1] [2] |
| *Pantoea sp.* MT58 | MT58 Wild type prototroph kanS | TEAM-3212 [3] |
| *Pantoea sp.* MT58 ΔIAI47_07670 | Strain with single gene deletion of IAI47_07670 | TEAM-3437 This study |
| *Pantoea sp.* MT58 ΔIAI47_09985 | Strain with single gene deletion of IAI47_09985 | TEAM-3423 This study |
| *Pantoea sp.* MT58 ΔIAI47_07670 ΔIAI47_09985 | Strain with double deletion of IAI47_07670 and IAI47_09985 | TEAM-3424 This study |
| *E. coli* WM3064 | *thrB1004 pro thi rpsL hsdS lacZΔM15 RP4-1360 Δ(araBAD)567 ΔdapA1341::[erm pir]* | [4] |

**Table S2.** Plasmids used in this study.

| Plasmid | Description | Reference |
|---|---|---|
| pTE1082 | Flanking 1 kb homology arms to generate ∆IAI47_09985 *pir+ R6K sacB kanR* | TEAM-3392 This study |
| pTE1086 | Flanking 1 kb homology arms to generate ∆IAI47_07670 *pir+ R6K sacB kanR* | TEAM-3393 This study |

**Table S3.** Genotyping primers.

| Primer Name | Primer sequence (5' to 3') | Target locus |
|---|---|---|
| AC452_09985_in_F | gcaccgtctgtcaggttgc | IAI47_09985 |
| AC453_09985_in_R | tccggcagctcacgc | IAI47_09985 |
| AC454_09985_out_F | ctcaccgattttttgcaacccc | IAI47_09985 |
| AC455_09985_out_R | ccgcttgagacctttgcacag | IAI47_09985 |
| AC460_07670_in_F | atgactgctcattccagccc | IAI47_07670 |
| AC461_07670_in_R | ttccggaaaatccgtctccac | IAI47_07670 |
| AC462_07670_out_F | aagcaggcgaaagcctttatgg | IAI47_07670 |
| AC463_07670_out_R | gaagagatctacgacgccatgc | IAI47_07670 |

**Table S4.** Reference sequences for plotting the phylogenetic protein tree

| Protein Name | Bacteria | Protein Accession |
|---|---|---|
| YfkO, Oxygen-insensitive NADPH nitroreductase | *Bacillus cereus* | WP_098774145.1 |
| YdgI, Nitroreductase family protein | *Bacillus cereus* | HDR4732676.1 |
| YfkO, NAD(P)H-dependent oxidoreductase | *Bacillus licheniformis* | MGG3356387.1 |
| YcnD, NADPH-dependent oxidoreductase | *Bacillus subtilis* | WP_088272169.1 |
| YdgI, Nitroreductase family protein | *Bacillus subtilis* | WP_160221377.1 |
| EF0404, Nitroreductase family protein | *Enterococcus faecalis* | WP_228180239.1 |
| EF1181, NADPH-dependent oxidoreductase | *Enterococcus faecalis* | WP_231435662.1 |
| NfsB, Oxygen-insensitive NAD(P)H nitroreductase | *Escherichia coli* | WP_432398827.1 |
| NfsB, Oxygen-insensitive NAD(P)H nitroreductase | *Klebsiella pneumoniae* | STV13824.1 |
| PnbA, Nitroreductase | *Lactiplantibacillus plantarum* | WP_319077754.1 |

| | | |
|---|---|---|
| Ntr, Nitroreductase | *Pantoea sp.* | CP040095.1 |
| Ntr1, Nitroreductase/dihydropteridine reductase | *Pantoea sp.* | CP024637.1 |
| NemA2, N-ethylmaleimide reductase2 | *Pantoea sp.* | CP082293.1 |
| NemA3, N-ethylmaleimide reductase3 | *Pantoea sp.* | CP060593.1 |
| AzoR2, FMN-dependent NADH-azoreductase AzoR2 | *Pseudomonas aeruginosa* | CEI20283.1 |
| AzoR3, FMN-dependent NADH-azoreductase AzoR3 | *Pseudomonas aeruginosa* | WP_049347706.1 |
| XenR_B, Xenobiotic reductase B | *Pseudomonas fluorescens* | AAF02539.1 |
| AzoA, Aerobic azoreductase | *Pigmentiphaga kullae* | AAO39146.1 |
| AzoB, NAD(P)-dependent oxidoreductase | *Pigmentiphaga kullae* | WP_130361747.1 |
| PnrB, Oxygen-insensitive NAD(P)H nitroreductase | *Pseudomonas putida* | WP_449432249.1 |
| XenR_A, Xenobiotic reductase XenA | *Pseudomonas putida* | WP_136916105.1 |
| SnrA, Oxygen-insensitive NADPH nitroreductase | *Salmonella typhimurium* | HAR9802468.1 |
| Frase1, NAD(P)H-dependent oxidoreductase | *(Alii)Vibrio fischeri* | GEK11990.1 |
| AzoB, FMN-dependent NADH-azoreductase | *Xenophilus azovorans* | WP_038202900.1 |
| Human Quinone reductase NQO1 (outgroup) | *Homo sapiens* | 1D4A_A |

**Table S5.** Residual TNT concentration (mg/L) in *P. putida* KT2440 cultures at 0 and 24 h. Note that the limit of detection from this standard curve is 0.78 mg/L, and values below that peak area are reported as 0 mg/L.

| Samples | TNT residual conc. (mg/L) |
|---|---|
| T0_TNT_1 | 20.8 |
| T0_TNT_2 | 21.58 |
| T0_TNT_3 | 19.1 |
| T0_TNT_4 | 21.1 |
| T24_TNT_1 | 0 |
| T24_TNT_2 | 0 |
| T24_TNT_3 | 0 |
| T24_TNT_4 | 0 |

**Table S6.** Proteomics data of **(A)** *P. putida* KT2440 and **(B)** *Pantoea sp.* MT58. The data are provided as a separate Excel file Supplementary Table S6, with the following headers: Protein.Group, Protein.Names, Protein, Protein.Description, Sample, Counts_mean, Counts_std, %Coeff_of_Variation(CV%), Counts_mean_sem, Sample_size, and Z-score.

**Table S7.** RB-TnSeq gene fitness scores for an oxidoreductase (PP_4931) and an azoreductase (PP_2866) under TNT conditions across four replicates.

| Gene | Condition | Fitness score |
|---|---|---|
| PP_2866 | Ammonium chloride and TNT (N) | -0.3 |
| | Ammonium chloride and TNT (N) | 0.1 |
| | Ammonium chloride and TNT (N) | -0.4 |
| | Ammonium chloride and TNT (N) | -0.2 |
| PP_4931 | Ammonium chloride and TNT (N) | N.D. |
| | Ammonium chloride and TNT (N) | N.D. |
| | Ammonium chloride and TNT (N) | N.D. |
| | Ammonium chloride and TNT (N) | N.D. |

### Supplementary Figures

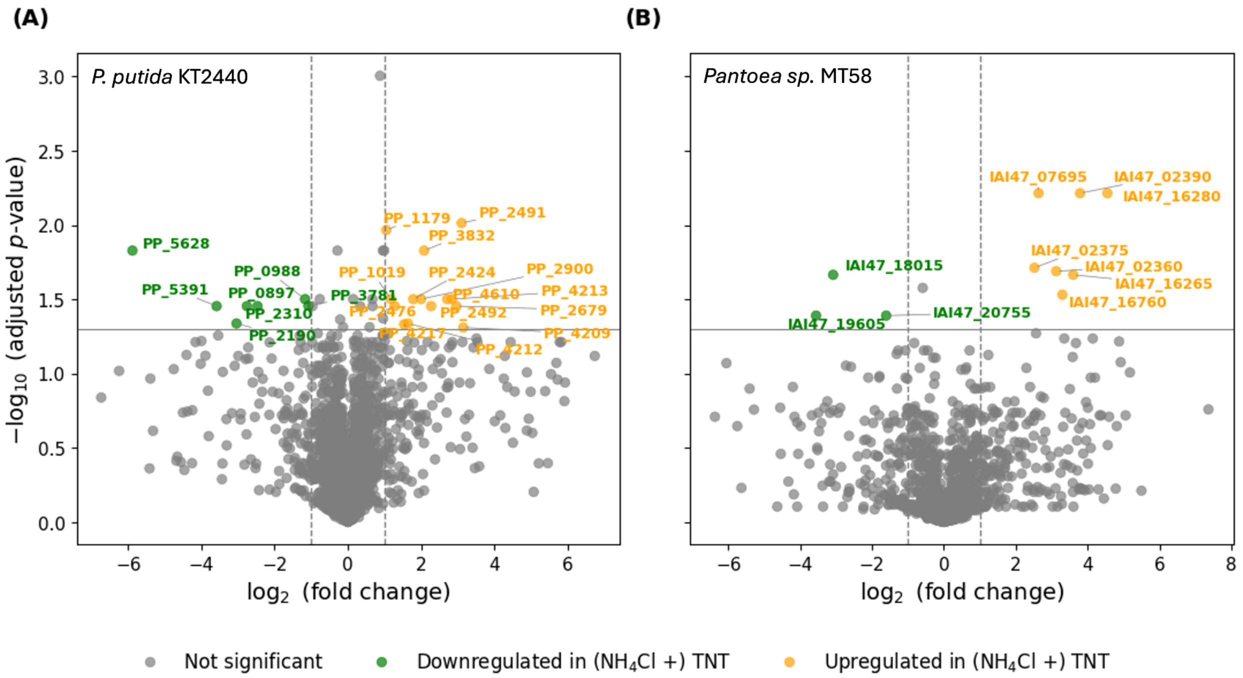


**Figure S1.** Adjusted *p*-value proteomics data of **(A)** *P. putida* KT2440 and **(B)** *Pantoea sp.* MT58, highlighting differentially expressed proteins when comparing growth in $NH_4Cl$ versus $NH_4Cl$ + TNT (KT2440) or TNT (MT58). The scatter plot shows $log_2$ fold change against the negative $log_{10}$ *p*-value for each protein, with proteins exhibiting a fold change of ±1 and an FDR-adjusted *p*-value < 0.05 highlighted in orange (upregulated) and green (downregulated) in the TNT condition.
